# SUPPA2 provides fast, accurate, and uncertainty-aware differential splicing analysis across multiple conditions

**DOI:** 10.1101/086876

**Authors:** Juan L. Trincado, Juan C. Entizne, Gerald Hysenaj, Babita Singh, Miha Skalic, David J. Elliott, Eduardo Eyras

## Abstract

Despite the many approaches to study differential splicing from RNA-seq, many challenges remain unsolved, including computing capacity and sequencing depth requirements. Here we present SUPPA2, a new method for differential splicing analysis that addresses these challenges and enables streamlined analysis across multiple conditions taking into account biological variability. Using experimental and simulated data SUPPA2 achieves higher accuracy compared to other methods; especially at low sequencing depth and short read length, with important implications for cost-effective use of RNA-seq for splicing; and was able to identify novel Transformer2-regulated exons. We further analyzed two differentiation series to support the applicability of SUPPA2 beyond binary comparisons. This identified clusters of alternative splicing events enriched in microexons induced during differentiation of bipolar neurons, and a cluster enriched in intron retention events that are present at late stages during erythroblast differentiation. Our data suggest that SUPPA2 is a valuable tool for the robust investigation of the biological complexity of alternative splicing.

## Introduction

Alternative splicing is related to a change in the relative abundance of transcript isoforms produced from the same gene (Lee and Rio, 2015). Multiple approaches have been proposed to study differential splicing from RNA sequencing (RNA-seq) data (Alamancos et al., 2014; Lahat and Grellscheid, 2016). These methods generally involve the analysis of either transcript isoforms (Froussios et al., 2017; Nowicka and Robinson, 2016; Sebestyen et al., 2015; Trapnell et al., 2013), clusters of splice-junctions (Hu et al., 2013; Vaquero-Garcia et al., 2016), alternative splicing events (Katz et al., 2010; Shen et al., 2014) or exonic regions (Anders et al., 2012). Relative abundances of the splicing events or transcript isoforms are generally described in terms of a percentage or proportion spliced-in (PSI) and differential splicing is given in terms of the difference of these relative abundances, or ΔPSI, between conditions (Venables et al., 2008; Wang et al., 2008). PSI values estimated from RNA-seq data have shown a good agreement with independent experimental measurements, and the magnitude of ΔPSI represents a good indicator of biological relevance (Katz et al., 2010; Venables et al., 2013). However, despite the multiple improvements achieved by recent RNA-seq analysis methods, many challenges remain unresolved. These include the limitations in processing time for current methods, the computational and storage capacity required, as well as the constraints in the number of sequencing reads needed to achieve high enough accuracy.

An additional challenge for RNA-seq analysis is the lack of robust methods to account for biological variability between replicates or to perform meaningful analyses of differentially splicing across multiple conditions. Although many methods assess the estimation uncertainty of the splicing event or transcript isoforms (Anders et al., 2012; Katz et al., 2010; Shen et al., 2014), they generally do so on individual events rather than considering the genome-wide distribution. Additionally, most methods determine the significance of differential splicing by performing tests directly on read counts, leaving the selection of relevant ΔPSI values to an arbitrary cut-off. In other cases, fold-changes instead of ΔPSI are given, which are even harder to interpret in terms of splicing changes.

We showed before that transcriptome quantification could be leveraged for the fast estimate of event PSI values with high accuracy comparing with experimental and simulated datasets (Alamancos et al., 2015). We now present here a new method for analyzing differential splicing, SUPPA2, which builds upon these principles to address the current challenges in the study of differential splicing, and taking into account biological variability. Compared with other existing approaches for differential splicing analysis using RNA-seq data, SUPPA2 provides several advantages. SUPPA2 can work with multiple replicates per condition and with multiple conditions. Additionally, SUPPA2 estimates the uncertainty of ΔPSI values as a function of the expression of transcripts involved in the event, taking into account all events genome-wide to test the significance of an observed ΔPSI, thereby directly estimating the biological relevance of the splicing change without relying on arbitrary ΔPSI cut-offs. Moreover, SUPPA2 incorporates the possibility to perform clustering of differentially spliced events across multiple conditions to identify groups of events with similar splicing patterns and common regulatory mechanisms. In conclusion, SUPPA2 enables cost-effective use of RNA-seq for the robust and streamlined analysis of differential splicing across multiple biological conditions. The software described here is available at https://github.com/comprna/SUPPA

## Results

### SUPPA2 monitors uncertainty to determine differential splicing

We showed before that the inclusion levels of alternative splicing events can be readily calculated from transcript abundances estimated from RNA-seq data with good agreement with experimental measurements and with other methods based on local measurements of splicing (Alamancos et al., 2015). SUPPA2 extends this principle to measure differential splicing between conditions by exploiting the variability between biological replicates to determine the uncertainty in the PSI values (see Methods). To illustrate our approach and to evaluate the dynamic range of SUPPA2 we used it to analyze RNA-seq data obtained after the double knockdown of TRA2A and TRA2B splicing regulators compared with controls (Best et al., 2014) (Fig. 1a). The differences in PSI value for each event between biological replicates are higher at low expression, in agreement with the expected higher variability at low read count. This biological variability provides information on the uncertainty of the PSI estimates. The significance of an observed ΔPSI value between conditions will depend on where in the distribution of the uncertainty falls. A large splicing change (|ΔPSI| value) may not be significant if it falls with a range of high uncertainty, whereas a small splicing change may be defined as robustly significant if it falls in the low uncertainty range. SUPPA2 estimates the significance considering the distribution between replicates for all events with similar transcript abundance; hence, it provides a lower bound for significant |ΔPSI| values that vary with the expression of the transcripts describing the event (Fig. 1b) (see Methods). The description of the uncertainty in terms of transcript abundances, given in transcript per million (TPM) units, rather than read counts provides several advantages. These include speed, as there is no need to store or go back to read information, as well as interpretability and application range, as transcript abundances are already normalized for transcript length and remain stable at different library sizes. More details on these advantages are provided below.

**Figure 1.**
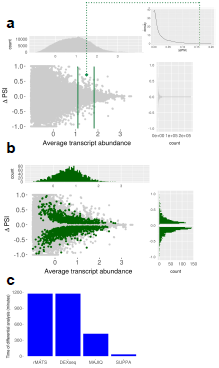
Overview of SUPPA2 differential splicing and time benchmarking analysis. (**a**)The central panel displays the ΔPSI values between replicates (*y*-axis) as a function of the average transcript abundance (*x*-axis), using data from (Best et al. 2015) (Methods). The attached panels display the ΔPSI values along the x-axis (top panel) and along the y-axis (right panel). The green dot represents an example of ΔPSI observed between conditions. The top-right panel shows the between-replicate |ΔPSI| density distribution against which an observed |ΔPSI| is compared to obtain a p-value. This density distribution is calculated from events with similar associated expression. (**b**) The central panel displays the ΔPSI values (y-axis) between conditions (green) or between replicates (gray) as a function of the average transcript abundance (x-axis) in log_10_(TPM+0.01) scale. Only events with p-value < 0.05 according to SUPPA2 are plotted in green. The attached panels display the distribution of the significant ΔPSI values along the x-axis (top panel) and along the y-axis (right panel). (c) Time performance of SUPPA2 compared to rMATS, MAJIQ and DEXSeq in the differential splicing analysis between two conditions, with 3 replicates each (Best et al. 2015). Time (y-axis) is given in minutes and in each case it does not include the read mapping, transcript quantification steps or the calculation of PSI values.

We compared SUPPA2 results with three other methods that calculate differential splicing using multiple replicates per condition: rMATS (Shen et al., 2014) and MAJIQ (Vaquero-Garcia et al., 2016), which describe changes in terms of ΔPSI, and DEXSeq (Anders et al., 2012), which uses fold-changes. Importantly, we found that SUPPA2 was much faster than the other methods, devoting 24 seconds to the PSI quantification and about 32 minutes and 47 seconds for differential splicing analysis on the same datasets (Fig. 1c). Since SUPPA2 performs the significance test directly on the ΔPSI values without needing to go back to the read data, it hence provides unmatched speed for differential splicing analysis. Comparing the results obtained with each method (Figure S1), we observed that rMATS and DEXSeq detect many apparently significant events with small inclusion changes that are not distinguishable from the variability between biological replicates, whereas SUPPA2 and MAJIQ separates well these two distributions. As SUPPA2 exploits the between-replicate variability to test for significance, it avoids the use of an arbitrary global |ΔPSI| threshold to identify biologically relevant events and detects significant events across a wide range of gene expression values (Figure S1). This feature of SUPPA2 should hence better rationalise |ΔPSI| threshold cut offs.

### SUPPA2 provides high accuracy at low sequencing depth and with short read lengths

To test the accuracy of SUPPA2 with different sequencing settings and compare it with other methods, we simulated 277 exon-cassette (SE) events and 318 alternative splice site (A5/A3) events with |ΔPSI|>0.2 between two conditions with 3 replicates per condition (Figure S2a). To perform a balanced comparison, we considered the same number of negative controls, consisting of different SE and A5/A3 events with arbitrary PSI values but with no simulated change between conditions (Table S1) (Methods). We simulated genome-wide RNA-seq reads using RSEM [18] at different sequencing depths: 120, 60, 25, 10 and 5 millions of 100nt paired-end reads per sample; and for different read-lengths: 100, 75, 50 and 25nt at a fixed depth of 25M paired-end reads. Despite the differences in the numbers and length of the reads (Table S2), the genes containing the positive and negative events used for benchmarking showed similar distributions of expression values at all depths and read lengths (Figure S2b). We then calculated differentially spliced events with SUPPA2, rMATS, MAJIQ and DEXSeq and evaluated the detection rate and accuracy on the simulated events (Table S3).

The detection rate was calculated as the proportion of simulated positive and negative cassette events that each method was able to measure from the RNA-seq data, i.e. the event was recovered regardless of whether it was detected as significant. The detection rate of SUPPA2 was superior than the other methods in all conditions, even at low depth and for shorter reads (Figure S2c). We also measured the true positives, i.e. the positive events that were observed to change significantly and in the same direction by each method; and the false positives, i.e. the negative events predicted to change significantly. For SE events the true positive rates were comparable across different sequencing depths (Fig. 2a). On the other hand, for shorter read length SUPPA2 recovered a higher proportion of true positives compared to the other methods (Fig. 2b). For A5/A3 events we also observed a similar decay in true positives with sequencing depth for all methods (Fig. 2c) and a higher accuracy of SUPPA2 with shorter read lengths (Fig. 2d). The same accuracies were observed if we imposed in addition the cutoff |ΔPSI|>0.2 for the predictions (Table S3). The reduced proportion of true positives at low depth and shorter read length in other methods was probably due to them relying on having sufficient junction and/or exonic reads. Additionally, even though SUPPA2 recovered in general more negative events, i.e. events simulated to be not differentially spliced; the false positive rate remained comparable to the other methods, and below 5% for all conditions (Table S3). To further evaluate the accuracies of the different methods, we computed Receiver Operating Characteristic (ROC) and Precision-Recall (PR) curves (Table S3). MAJIQ and SUPPA2 show similar areas under the ROC and PR curves, which drop at low depth and with short read lengths; whereas DEXSeq and rMATS lower areas across all values of depth and read length.

**Figure 2.**
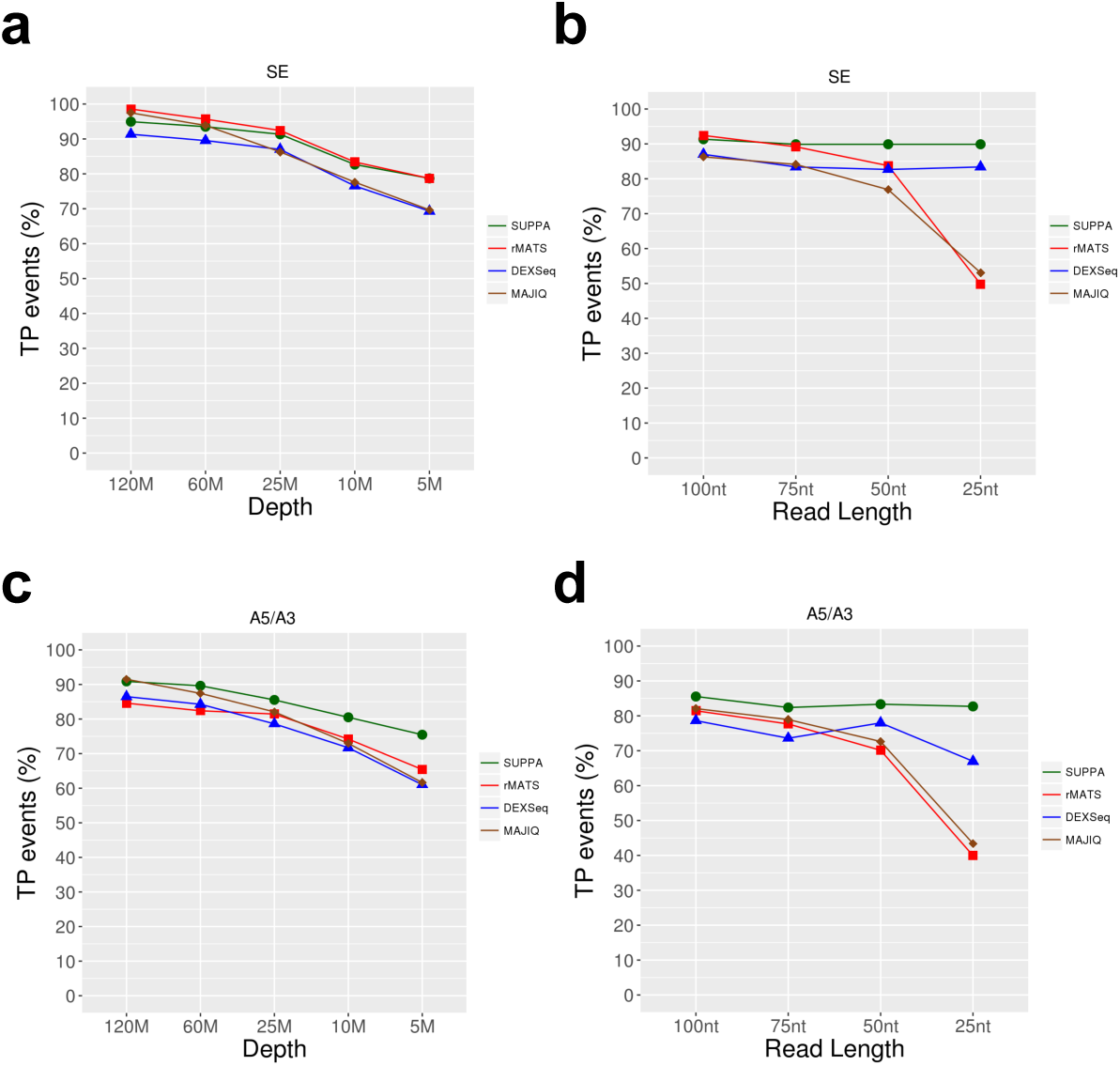
Accuracy analysis with simulated data. **(a)** Proportion of events measured by each method (y-axis) from the 277 positive simulated cassette events at different sequencing depths (x-axis), from 120 million (120M) down to 5 million (5M) of paired end reads, using 100nt paired-end reads. **(b)** As in (a) but for different read lengths (x-axis) at fixed depth (25M). **(c)** True positive (TP) rate (in terms of percentage) for each method (*y*-axis) at different sequencing depths (x-axis) for 100nt paired-end reads. TPs were calculated as the number of statistically significant events according to each method: corrected p-value<0.05 for SUPPA2, rMATS and DEXSeq; and posterior(|ΔPSI|>0.1)>0.95 for MAJIQ. **(d)** As in (c) but for different read lengths (x-axis) at fixed depth (25M).

We also considered an unbalanced configuration where one replicate had 120M reads and the other two replicates had 10M reads. In this hybrid configuration, SUPPA2 recovered a high number of events and high number of true positives for SE events. On the other hand, for A5/A3 events we observed a slight drop in accuracy (Table S3), probably due to a high proportion of short variable regions in the alternative sites events (79 events (25%) of the A5/A3 events involved a region of under 9 nucleotides), which may be more problematic for correct transcript quantification than using direct mapping to splice junctions. Importantly, although MAJIQ showed a high detection rate and accuracy in the unbalanced configuration, it had to be run with specialized parameters (Methods), whereas SUPPA2 was run in the same way for all cases. Additionally, SUPPA2 also showed high correlation values between the predicted and simulated ΔPSI values (Table S3), and similar to those obtained with rMATS and MAJIQ. In the light of these results, we can conclude that SUPPA2 performs comparably to other methods under a wide spectrum of sequencing conditions and in particular, it outperforms other methods at low sequencing depth and short read length.

### SUPPA2 provides accurate splicing change quantification compared with experimental results

To further evaluate the accuracy of SUPPA2 in recovering ΔPSI values we used 83 events that had been validated experimentally by RT-PCR upon TRA2A and TRA2B knockdown compared to control cells (Table S4) (Methods) (Best et al., 2014). For each method, we compared the ΔPSI estimated from RNA-seq with the ΔPSI from RT-PCR. SUPPA2 agreement to the RT-PCR ΔPSI values was similar to rMATS and MAJIQ (Fig. 3a) (Table S5). Using two other independent RT-PCR datasets published previously (Vaquero-Garcia et al., 2016), SUPPA2 also showed similar accuracy compared to rMATS and MAJIQ (Figures S3a and S3b) (Tables S6-S9). Finally, using 44 RT-PCR negative cassette events that did not show any significant change upon the double knockdown of TRA2A and TRA2B, SUPPA2 had a lower false positive rate compared to the other methods (Fig. 3b) (Tables S10-S11).

**Figure 3.**
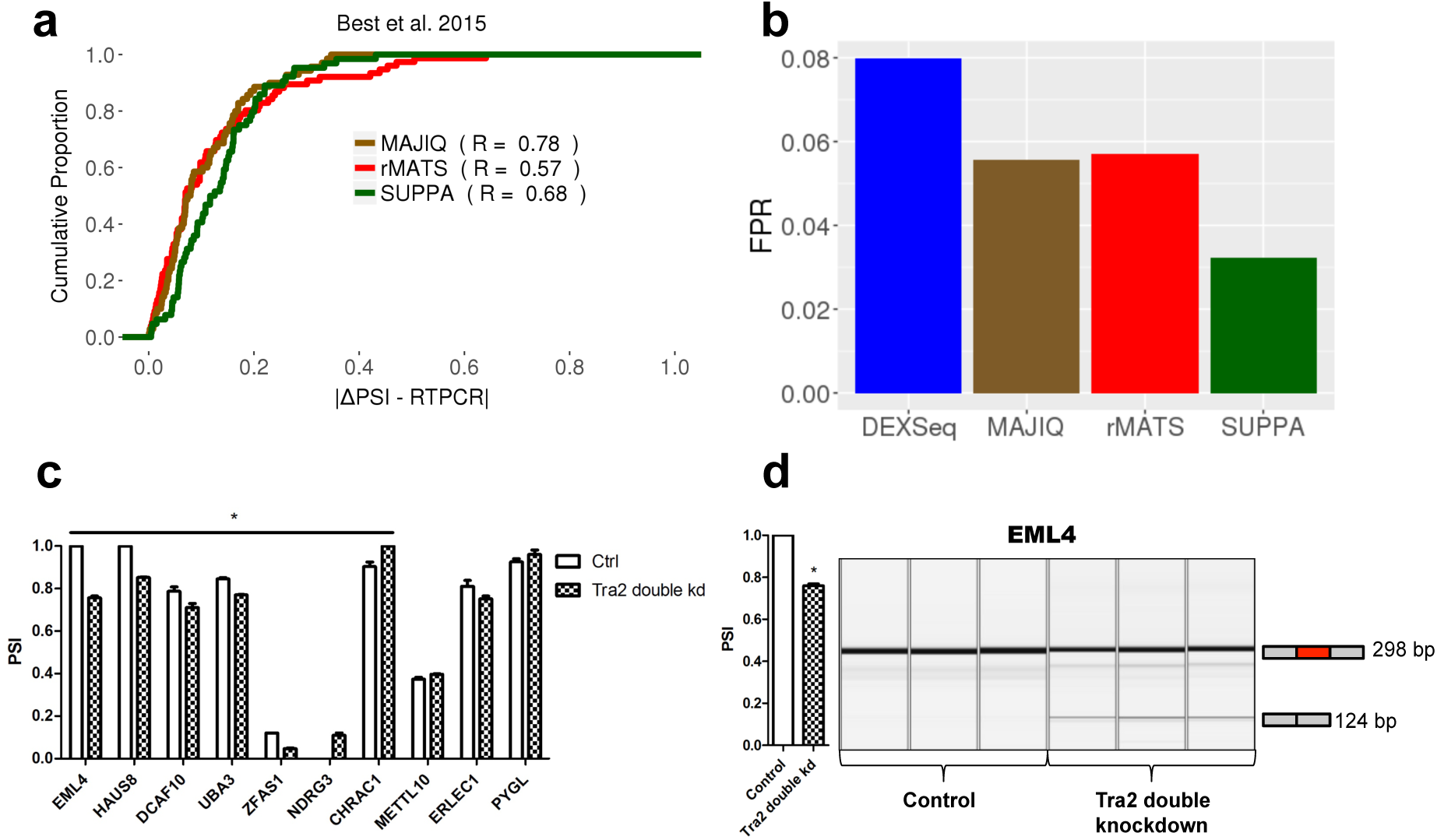
Experimental validation of differentially splicing predictions by SUPPA2. **(a)** Comparison of predicted and experimentally validated ΔPSI values for 83 cassette events differentially spliced between the double knockdown of TRA2A and TRA2B and control in MDA-MB-231 cells. We show the cumulative proportion of cases (y-axis) according to the absolute difference between the predicted and the experimental value (|ΔPSI - RTPCR|), for the events detected by each method: SUPPA2 (66), rMATS (78), and MAJIQ (72). Additionally, we give for each method the Pearson correlation R between predicted and experimental values, **(b)** False positive rate (FPR) calculated using 44 RT-PCR negative events. FPR was calculated as the proportion of the detected events that was found as significant by each method: SUPPA2 (1/31), rMATS (2/35), MAJIQ (2/36), DEXSeq(2/25). **(c)** Experimental validation by RT-PCR of a subset of novel events with TRA2B CLIP tags and Tra2 motifs. These events include cases that were only predicted by SUPPA2 (CHRAC1, NDRG3, METTL10), and cases that were not predicted by any method but were significant according to SUPPA2 before multiple test correction (ERLEC1, PYGL, DCAF10, HAUS8, EML4, UBA3) (Table S14). RT-PCR validation was performed in triplicate. Error bars indicate the standard error of the mean. Cases that change significantly (p<0.05) according to a two-tailed t-test comparing the three values of the knockdown versus control are indicated with an asterisk. **(d)** Experimental validation of a new skipping event in *EML4* upon knockdown of TRA2A and TRA2B (3 biological replicates shown in each case).

### SUPPA2 identifies experimentally reproducible splicing changes not detected by other methods

The results described above suggest a general agreement between the different methods in the detection of significant differentially spliced events. To assess this question, we performed a direct comparison of the results obtained from the four methods SUPPA2, rMATS, MAJIQ and DEXSeq, using the same RNA-seq data for the knockdown of TRA2A and TRA2B compared with controls [17]. Since exon-cassette (SE) (48.71%) and alternative splice-site (A5/A3) (37.71%) events are the most frequent events in humans compared to mutual exclusion (6.22%) or intron-retention (7.36%), we decided to match SE and A5/A3 events across all four methods. We were able to identify 7116 SE events and 2924 A5/A3 events unambiguously detected by all four methods, i.e. they were measured and tested for significance by all methods (Figure S4a) (Table S12) (Methods).

For the 7116 SE events, each method found between 133 and 274 events to be significant, with 370 events predicted as significant by any one method, but only 22 events predicted by all four methods (Figure S4a). Similarly, 352 A5/A3 events were predicted to be significant by at least one method, and only 2 predicted by all four methods (Figure S4a). Events detected by more methods tended to have higher ΔPSI values (Figure S4b) and covered a smaller range of gene expression values (Figure S4c). Despite the low detection overlap, the significant events predicted by each method independently showed enrichment of TRA2B CLIP tags and of Tra2 binding motifs (Table S13) (Supplementary Methods); hence, each set independently had the expected properties related to the knockdown experiment. It is possible that each method describes a different subset of changes and generally misses others. To seek further support for this point, we selected for experimental validation 15 SE events and 7 A3 events that had CLIP tags and Tra2 motifs nearby the regulated exon. The 7 A3 events and 6 of the 15 SE events were predicted only by SUPPA2, whereas the remaining 9 were not predicted by any of the four methods, but were significant according to SUPPA2 before multiple test correction (Table S14). From these 15 SE events, 5 of them only showed one PCR band and could not be evaluated. However, for the rest, 7 changed significantly according to the RT-PCR (2-tailed t-test p-value<0.05), with 6 of them changing in the same direction predicted by SUPPA2. Overall, 9 events changed in the same direction as predicted (Fig. 3c) (Table S14). In particular, we validated a new event in *EML4* (Fig. 3d), a gene involved in cancer through a fusion with *ALK* that is not present in MDA-MB-231 cells (Lin et al., 2009). In addition, we could measure 6 of the 7 A3 events; all were measured to change in the same direction as predicted by SUPPA2 and 4 were significant (2-tailed t-test p-value<0.05) (Table S14). This analysis shows the value of using a suite of methods based on different algorithms, like SUPPA2, to reveal novel experimentally reproducible events that are missed by other methods.

### SUPPA2 finds biologically relevant event clusters across multiple conditions

SUPPA2 is also able to analyze multiple conditions by computing the pairwise differential splicing between conditions, and can detect groups of events with similar splicing patterns across conditions using density-based clustering (Methods). To evaluate the ability of SUPPA2 to cluster events, we analyzed a 4-day time-course of differentiation of human iPSCs into bipolar neurons (Busskamp et al., 2014), which had not been analyzed yet for alternative splicing. SUPPA2 identified 2780 regulated cassette events (p-value < 0.05), out of which 207 (8,4%) were microexons (length < 28nt), which represent an enrichment (Fisher’s exact test p-value < 2.2e-16, odds-ratio= 3.94) compared to a set of 20452 non-regulated cassette events (p-value > 0.1), with the majority of these microexons (69%) significantly more included in differentiated cells (ΔPSI > 0 and p-value < 0.05 between the first and fourth day).

We evaluated the performance of the two density-based cluster methods implemented in SUPPA2, DBSCAN (Ester et al., 1996) and OPTICS (Ankerst et al., 1999), using different input parameters. In spite of OPTICS requiring more computing time than DBSCAN (43s vs 5s), it produced slightly better clustering results (Figures S5a-S5d) (Table S15). For a maximum reachability distance of 0.11, i.e. maximum distance of an event to a cluster to be considered part of the cluster, we obtained 3 well-differentiated clusters (silhouette score = 0.572) (Figs. 4a-c) (Table S16). Cluster 0 increased inclusion at late steps of differentiation and showed an enrichment in microexons (32 out of 115 events) with respect to unclustered regulated cassette events (Fisher’s exact test p-value = 0.0148, odds-ratio = 5.3521). In contrast, clusters 1 and 2 decreased inclusion with differentiation, and contained 2 (out of 20 events) and no microexons, respectively. These results are in agreement with the previously observed enrichment of microexon inclusion in differentiated neurons (Irimia et al., 2014; Li et al., 2015).

**Figure 4.**
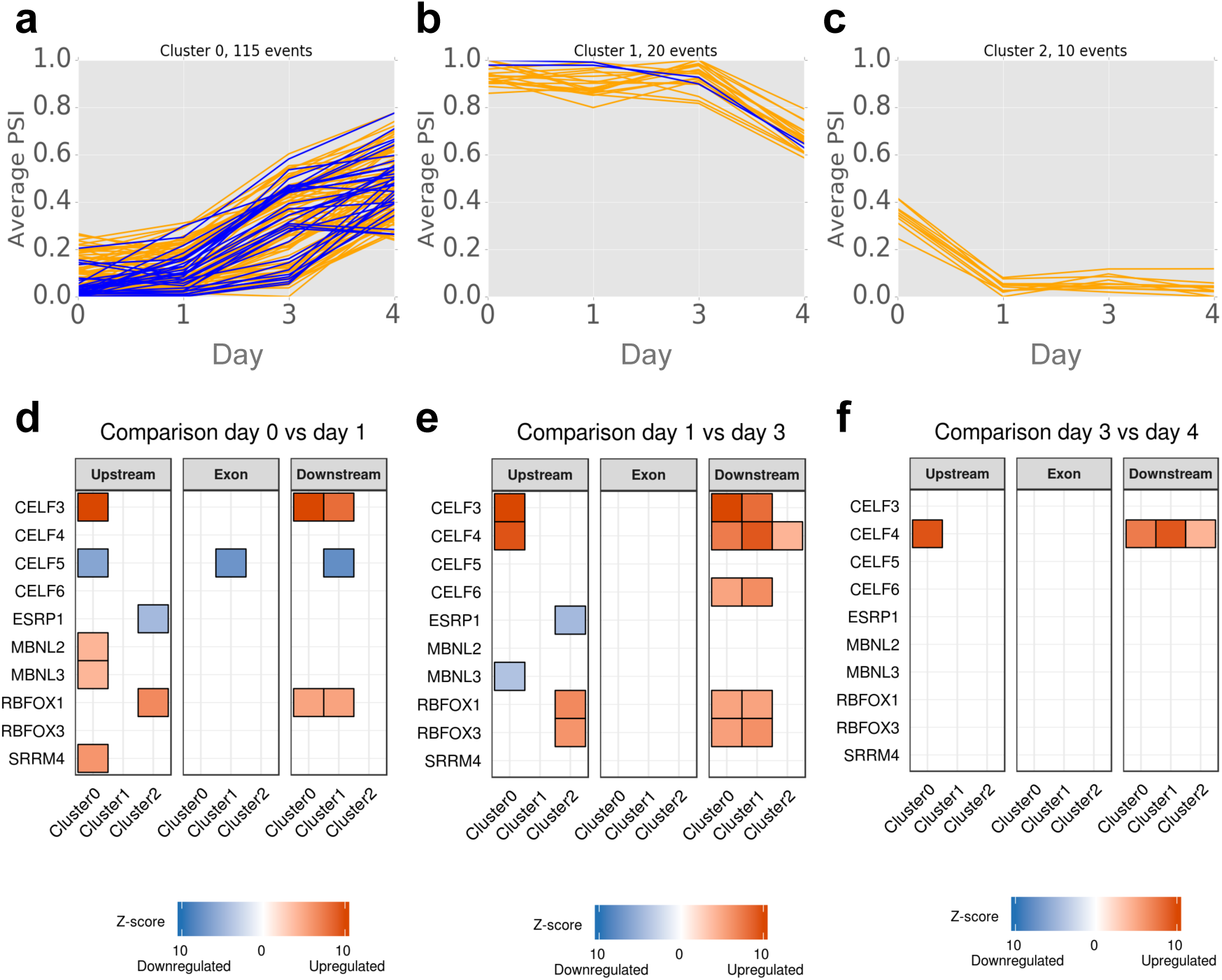
Prediction and clustering of differentially spliced events across bipolar neuron differentiation. Density-based clustering performed on the 2780 regulated cassette events that change splicing significantly in at least one comparison between adjacent steps across 4 differentiation stages (days after differentiation 0,1,3,4). Plots **(a-c)** show the average PSI (*y*-axis) per stage (*x*-axis) of the events in the three clusters obtained. Microexons (<28nt) are plotted in blue over the rest of the events in orange. **(d-f)** Motif enrichment associated to each of the 3 clusters in (a-c) in the regions upstream (200nt), exonic, and downstream (200nt). Only enriched motifs associated to splicing factors that are differentially expressed are shown in each comparison between differentiation stages (days after differentiation 0,1,3,4). In red we indicate the splicing factors that are upregulated and in blue those that are downregulated at each stage. The color intensity indicates the z-score of the motif enrichment. Motifs are shown in each cluster and region where they are found enriched.

To further validate the findings with SUPPA2, we performed a motif enrichment analysis in regulated events compared to non-regulated events. Notably, compared to the non-regulated events, the 2780 regulated cassette events showed enrichment in binding motifs for the RNA binding protein (RBP) SFPQ (z-score > 4), which has been described before as a necessary factor for neuronal development (Lowery et al., 2007). Additionally, the differentially spliced events in clusters were enriched in, among others, *CELF, RBFOX, ESRP, MBNL and SRRM4* motifs (Fig. 4d-4f), in concordance with the described role of *CELF, RBFOX* and *SRRM4* genes in neuronal differentiation (Kim et al., 2013b; Li et al., 2015; Norris et al., 2014; Raj et al., 2014). Consistent with these findings, *SRRM4* and members of the *CELF* and *RBFOX* families showed upregulation at the initial steps of iPSC differentiation into neurons (Figure S5) (Table S17). On the other hand, *CELF5* and *ESRP1* were downregulated during differentiation. The *MBNL3* gene showed initial upregulation at stage 1, followed by downregulation at later stages (Figure S5) (Table S17). Notably, we found that only the cluster enriched in microexon splicing inclusion showed an enrichment of SRRM4 motifs upstream of the regulated exons, in agreement with the previous description of SRRM4 binding upstream of microexons to regulate their inclusion during neuronal differentiation (Raj et al., 2014), and further supports the specificity of SRRM4 to regulate microexons. Our results also suggest possible novel regulators of neuronal differentiation, such as the *MBNL* proteins in the regulation of events increasing exon inclusion and *ESRP* in events that decrease exon inclusion (Fig. 4d-4f).

We also used SUPPA2 to analyze differential splicing across 5 stages of erythroblast differentiation (Pimentel et al., 2014). In this case we considered all event types for clustering. For the optimal value of maximum reachability distance (S=0.1), we obtained two homogeneous and well-differentiated clusters (silhouette score = 0.91), one for events with low PSI that increased at the last differentiation stage with 149 events, and a second cluster with 86 events that showed the opposite behaviour (Figure S6). In agreement with previous results (Pimentel et al., 2016), we observed an enrichment of intron retention events in the cluster of events that increased inclusion at the late differentiation stage, as compared with the other cluster, which does not include any retained intron (Fisher’s exact test p-value = 0.04958). We conclude that SUPPA2 provides a powerful approach to analyze splicing across multiple conditions, validated not only by intrinsic measures of clustering consistency, but also by recovering known biological results and new features.

## Discussion

Our extensive evaluations here indicated that SUPPA2 provides a broadly applicable solution to current challenges in the analysis of differential splicing from RNA sequencing data across multiple conditions, and has features that will make it attractive to many potential users. SUPPA2 is faster than other methods, and maintains a high accuracy, especially at low sequencing depth and for short read-length. Despite using less reads or shorter reads, SUPPA2 could detect the majority of the simulated events and maintained a high proportion of true positives and low proportion of false positives. SUPPA2 thus offers an unprecedented opportunity to study splicing in projects with limited budget, or to reuse for splicing studies available sequencing datasets with lower depth than usually required by other methods. Additionally, the low computing and storage requirements of SUPPA2 makes possible to perform fast differential splicing processing and clustering analysis on a laptop. Thus, coupled with fast methods for transcript quantification (Bray et al., 2016; Patro et al., 2014, 2017), SUPPA2 facilitates the study of alternative splicing across multiple conditions without the need for large computational resources. The simplicity and modular architecture of SUPPA2 also makes it a very convenient tool in multiple contexts, as PSI values from other methods and for other event types, like complex events, or data types, like transcripts, can be used in SUPPA2 for differential splicing analysis or for clustering across conditions.

According to our simulated benchmarking analysis, as well as others published before, it may seem that bioinformatics methods used to analyze RNA-seq data tend to coincide on a large number of events. However, using real experimental data we actually observed low agreement in targets between methods. These discrepancies in target selection can be explained by various factors, including the different ways in which a splicing change is represented by each method (e.g. an event, an exon or a graph), how changes in splicing patterns are tested by each method, and how biological and experimental variability affects these tests. Intriguingly, the results from each method do make sense biologically, in that differentially spliced events were enriched in motifs and mapped protein-RNA interaction sites related to the depleted splicing factor. This makes it unlikely that any one method provides a clear advantage in terms of the results, and instead suggests that at least two or three methods should be used to identify all the possible significant splicing variants between different conditions. In particular, we chose for comparison three other methods with very different representations of the splicing and statistical approach. The results we obtained recommend use of two or more such tools to comprehensively monitor splicing complexity by picking out different sets of events that would not otherwise be discovered, rather than identifying largely overlapping groups of events. Supporting this point we could validate experimentally events not predicted by any other methods but predicted by SUPPA2. We further observed that although most methods had the power to identify small significant ΔPSI values, different methods tended to agree on events with large splicing changes. Importantly, a fraction of these significant events with small ΔPSI are indistinguishable from the variability observed between replicates and hence are not likely to be biologically relevant. SUPPA2 also performs a statistical test that can separate significant splicing changes from the biological variability, providing thus an advantage to identify biologically relevant changes across wide range of expression values. By exploiting the biological variability, without having to go back to the read data, SUPPA2 provides a fast and accurate way to detect differential splicing without the need for arbitrary global ΔPSI thresholds.

Although SUPPA2 relies on genome annotation to define events, poorly annotated genomes can be improved and extended before analysis by SUPPA2. In fact, recent analyses have shown that improved annotations lead to significantly better PSI estimates from RNA-seq when benchmarked to high-resolution RT-PCR measurements (Brown et al., 2016; Zhang et al., 2015, 2017). Current technological trends predict an increase in the number of efforts to improve the transcriptome annotation in multiple species and conditions (Garalde et al., 2016). In this direction, SUPPA2 could play a key role for the systematic and rapid genome-wide analysis of splicing following annotation and sample updates.

## Conclusions

In conclusion, the speed, modularity and accuracy of SUPPA2 enable cost-effective use of RNA sequencing for the robust and streamlined analysis of differential splicing across multiple biological conditions.

## Methods

### Differential splicing

SUPPA2 uses transcript quantification to compute inclusion values (PSI) of alternative splicing events across multiple samples. Given the calculated PSI values per sample, SUPPA2 considers two distributions: one for the ΔPSI values between biological replicates and one for the ΔPSI values between conditions. For the first distribution, for each event SUPPA2 calculates the ΔPSI value between each pair of biological replicates together with the average abundance of the transcripts describing the event across the same replicates:

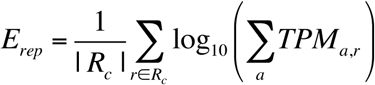

where *r*=*1*,.,|*R_c_*| runs over the replicates in each condition *c*=*1,2,* and *a* indicates the two or more transcripts describing the event, and *TPM_a,r_* indicates the abundance of transcript *a* in replicate *r* in transcript per million (TPM) units. For the distribution between conditions, the ΔPSI values are calculated as the difference of the means in the two conditions, together with the average abundance of transcripts describing the event across both conditions for each event:

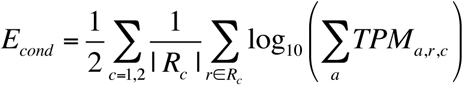

where *TPM_a,r,c_* indicates the abundance of transcript *a* in replicate *r* in condition *c* in transcript per million (TPM) units. Given the observed ΔPSI and *E_cond_* values for an event between conditions, its significance is calculated from the comparison with the ΔPSI distribution between replicates for events with *E_rep_* values in the neighborhood of the observed *E_cond_*. This neighborhood is defined by first selecting the closest value *E*_rep_* from all points *i* from the between-replicate distribution:

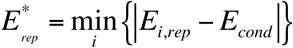

using binary search and selecting a fixed number of events (1000 by default) around the *E*_rep_* value in the interval or ordered values. The selected events define an empirical cumulative density function (ECDF) over |ΔPSI| from which a p-value is calculated:

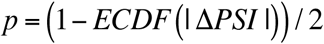

Here we implicitly assume that the background distribution is symmetric. SUPPA2 includes an option to correct for multiple testing using the Benjamini-Hochberg method across all events from the same gene, as they cannot be considered to be entirely independent of each other, for which the false discovery rate (FDR) cut-off can be given as input.

### Clustering

SUPPA2 currently implements two density-based clustering methods: DBSCAN (Ester et al., 1996) and OPTICS (Ankerst et al., 1999). Density-based clustering has the advantage that one does not need to specify the expected number of clusters, and the choice between the two methods depends mainly on the computational resources and the amount of data. Both methods use the vectors of mean PSI values per event and require as input the minimum number of events in a cluster (N), which could be interpreted as the minimum expected size of the regulatory modules. OPTICS also requires as the maximum reachability distance (S), which represents the maximum distance in PSI space of an event to a cluster. On the other hand, DBSCAN requires as input the maximum distance to consider two events as cluster partners (D), which OPTICS calculates through an optimization procedure allowing any value below S.

DBSCAN allows to perform simple and fast data partitioning but has the drawback of being sensitive to the input parameters. On the other hand, OPTICS, which can be seen as a generalization of DBSCAN, explores the possible maximum values for D beyond which clustering quality drops. OPTICS can thus potentially produce better clustering results, since it is not limited to a fixed radius of clustering, but it is penalized by a greater computational cost. Clustering is performed only with events that change significantly in at least one pair of adjacent conditions. Three different distance metrics can be currently used: Euclidean, Manhattan and Cosine. Cluster qualities are reported using the silhouette score (Rousseeuw, 1987), which indicates how well the events are assigned to clusters; and the root mean square standard deviation (RMSSTD), which measures the homogeneity of each cluster. Additionally, the number and percentage of events in clusters is also reported. Motif enrichment analysis was performed as before (Sebestyén et al., 2016) using MOSEA, available at https://github.com/comprna/MOSEA. Further details on the motif enrichment and analysis of differential expression are provided in Supplementary Material.

### Simulated datasets

For the simulation we used the quantification of RefSeq transcripts for the 3 control samples from (Best et al., 2014) (GSE59335) with Salmon (Patro et al., 2017) as theoretical abundances, and considered genes with only two isoforms containing an skipping exon (SE) or alternative splice site (A5/A3) event and only 1 associated event. For the benchmarking analysis, we selected a set of positive and a set of negative events for each event type with the same number of randomly chosen events, 277 for SE events and 318 for A5/A3 events. For the positive set we simulated differential splicing by exchanging the theoretical abundance of their associated transcripts values. We selected to be positive events only those having an absolute difference of relative abundance greater than 0.2, so that the simulated change was sufficiently large:

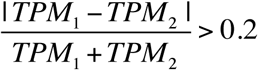

where TPM1 and TPM2 are the abundances for the two transcripts in the gene, given in transcript per million (TPM) units. For the negative set, we took an equal number of events without exchanging their TPM values. These negative events had a gene expression distribution and a distribution of transcript relative abundance similar to the positive events, and an expected variability between conditions similar to the variability between biological replicates.

We used RSEM (Li and Dewey, 2011) to simulate sequencing reads for the 2 conditions, 3 replicates each, at various depths: 120, 60, 25, 10 and 5 millions of 100nt paired-end reads per sample, and at various read lengths: 100nt, 75nt, 50nt and 25nt, at depth of 25M paired-end reads (Tables S1-S3). Further details of the simulations are given in the Additional file 3:Supplementary Material. Datasets and commands to reproduce these simulations are available at https://github.com/comprna/SUPPA_supplementary_data.

### Experimental datasets

We analyzed RNA-seq data for the double knockdown of TRA2A and TRA2B in MDA-MB-231 cells and controls with 3 replicates per condition (Best et al., 2014) (GSE59335). For benchmarking, we used 83 RT-PCR validated events for comparison (Tables S4-S5) and 44 RT-PCR negative events (Tables S12-S13). We also analyzed data from Cerebellum and Liver mouse tissues covering 8 different time points from 2 full circadian cycles (Zhang et al., 2014) (GSE54651) and performed a comparison with 50 events validated by RT-PCR (Vaquero-Garcia et al., 2016) comparing samples CT28, CT40 and CT52 in Cerebellum with the same circadian time points in Liver (Tables S8-S9). We also analyzed RNA-seq data for stimulated and unstimulated Jurkat T-cells and compared it with RT-PCR validated events (no tested replicates) (Cole et al., 2015; Vaquero-Garcia et al., 2016) (SRP059357) (Tables S10-S11). From these 54 RT-PCR validated events, we only used the 30 events that had experimental value |ΔPSI|>0.05. For the study of multiple conditions, we used RNA-seq samples from a 4-day time-course for the differentiation of human iPSCs cells into bipolar neurons (Busskamp et al., 2014) (GSE60548). Original data was for days 0,1,3,4 after initiation of differentiation. Additionally, we analysed RNA-seq from 5 steps of differentiating human erythroblasts (Pimentel et al., 2016) (GSE53635), with 3 replicates per condition. RNA-seq reads from all experiments were used to quantify human and mouse transcripts from Ensembl (version 75 - without pseudogenes) with Salmon (Patro et al., 2017). Reads were mapped to the human (hg19) or mouse (mm10) genomes using TopHat (Kim et al., 2013a). All methods other than SUPPA2 were used with these mappings. Cassette events from SUPPA2 and rMATS were matched to the RT-PCR validated events in each dataset, considering only those cases where the middle exon matched exactly the validated exons and confirming the flanking exons with the RT-PCR primers when available. Ambiguous matches were discarded from the comparison. For MAJIQ we selected the inclusion junction compatible with the validated event that had the largest posterior probability for |ΔPSI|>0.1. For DEXSeq we considered only exonic regions that matched exactly with the regulated exon of the experimentally validated cassette event. To select a set of cassette events common to all four methods, we selected the events measured by both SUPPA2 and rMATS such that the middle exon matched exactly a DEXSeq exonic region and did not appear in more than one event from SUPPA2 or rMATS. From this set, we selected those for which any of the two inclusion junctions was present in MAJIQ, and selected the junction with the largest posterior probability for |ΔPSI|>0.1. Further details are provided in Supplementary Material.

### Time performance

Running time was measured using the Unix time command *time*. For SUPPA2 running time was measured independently of the transcript quantification step. Similarly, for all other methods the running time did not include the read-mapping step. Time was measured independently for PSI calculation and for differential splicing analysis. All methods were run on a Unix machine with 12Gb of RAM and 8 Intel Xeon 2GHz CPU cores.

### Experimental validation

Details on the experimental validation are given in Supplementary Material.

### Software and datasets

SUPPA2 is available at https://github.com/comprna/SUPPA Commands and datasets used in this work are available at https://github.com/comprna/SUPPA_supplementary_data Software for the motif enrichment analysis is available at https://github.com/comprna/MOSEA

## List of abbreviations

PSI: proportion spliced in, RT-PCR: reverse transcriptase polymerase chain reaction, iPSC: induced pluripotent stem cell, CLIP: cross linking immunoprecipitation, TRA2A/B: Transformer-2 protein homolog alpha/beta, RNA-seq: RNA sequencing, TPM: transcripts per million.

## Acknowledgements

The authors thank C. Calixto, J. Brown, R. Zhang, M. Irimia, and N. Barbosa-Morais, for useful discussions and to J. Vaquero-Garcia and Y. Barash for comments on an earlier version of the manuscript. This work was supported by the MINECO and FEDER (BIO2014-52566-R) and AGAUR (SGR2014-1121), the BBSRC (BB/P006612/1), and Breast Cancer Now (2014NovPR355). GH is a BBSRC-funded PhD student.

## Author Contributions

J.C.E, M.S. and EE designed and implemented the method and algorithms, J.C.E., J.L.T., and E.E. devised the analyses and J.C.E, J.L.T, and M.S. carried out the benchmarking analyses. G.H and D.J.E. produced the datasets related to the double knockdown of TRA2A and TRA2B and performed the validation experiments. B.S. carried out the software development and analysis for motif enrichment analyses. J.C.E, J.L.T. and EE wrote the manuscript with essential input from G.H. and D.J.E.

